# How much can reticulate evolution entangle plant systematics? Revisiting subfamilial classification of the Malvatheca clade (Malvaceae) on the basis of phylogenomics

**DOI:** 10.1101/2025.10.17.683170

**Authors:** Gustavo Luna, Lucas Costa, Flávia Fonseca Pezzini, Nisa Karimi, Joeri Sergej Strijk, Jefferson Guedes Carvalho-Sobrinho, Matheus Colli-Silva, André Marques, Gustavo Souza

## Abstract

Reticulate evolution (RE), involving hybridization and related processes, generates network-like rather than strictly bifurcating relationships among lineages and can obscure phylogenetic relationships. Detecting ancient hybridization is particularly challenging, as genomic signals may erode over time. The Malvatheca clade (Malvaceae), marked by multiple paleopolyploidy events since it’s estimated origin 66 my, offers a useful model for examining RE. Its three subfamilies—Bombacoideae (with high chromosome numbers, mostly trees), Malvoideae (lower chromosome numbers, mostly herbs), and the recently described Matisioideae—show unresolved relationships, with several taxa of uncertain placement. We conducted a phylogenomic analysis of 62 Malvatheca species via complete plastomes, 35S rDNA cistrons, and comparative repeatome data. All datasets consistently resolved three main clades: (i) Bombacoideae, (ii) Malvoideae, and (iii) a heterogeneous assemblage including most of the recently assigned Matisioideae and several *incertae sedis* taxa, suggesting a more complex evolutionary history and circumscription than previously recognized. Chromosome numbers were negatively correlated with repeatome diversity: Bombacoideae presented higher counts but lower repeat diversity, possibly reflecting slower repeat evolution associated with woody growth forms. In contrast, clade III showed marked heterogeneity in both chromosome number and repeat composition, which is consistent with a reticulate origin. Overall, our results show evidence of ancient hybridization and polyploidy, predominantly allopolyploidy, in shaping Malvatheca evolution. These results highlight that reticulation and genome dynamics, rather than taxonomic boundaries alone, are central to understanding the diversification of Malvatheca.

## Introduction

Reticulate evolution (RE) refers to processes in which genetic material is exchanged between lineages, producing network-like rather than strictly bifurcating phylogenetic relationships (Arnold, 1997; Gontier, 2015). It can arise through hybridization, horizontal gene transfer, or endosymbiosis, and challenges the traditional tree-like view of evolution (Mallet, 2007). Reticulation also complicates classification by blurring taxonomic boundaries and generating conflicting phylogenetic signals. In plants, where hybridization is common, genetic and phenotypic distinctions may be obscured, leading to uncertain or misleading taxonomic boundaries (Mallet, 2007).

Polyploidy (whole genome duplication, WGD), the presence of multiple complete genomes in a species, is closely linked to reticulation and is a recurrent feature of angiosperm evolution (Bowers & Paterson, 2021). Allopolyploidization, i.e., hybridization followed by genome doubling, has been recognized as a major driver of plant diversification (Paterson et al., 2004; Jiao et al., 2011; Vekemans et al., 2012). Notably, a burst of polyploidization events in flowering plants occurred near the Cretaceous–Paleogene (K–Pg) boundary (∼66 Mya), coinciding with a mass extinction event (Vanneste et al., 2014; Fawcett & Van de Peer, 2010). The clustering of independent WGDs across angiosperms during this period suggests that polyploidy promoted survival and diversification by providing raw material for genetic novelty (Fawcett et al., 2009; Wood et al., 2009).

Studying RE in ancient lineages remains difficult because signals of hybridization and polyploidy are often eroded over time (Zhang et al., 2025). Additionally, disentangling processes such as incomplete lineage sorting, horizontal transfer, or genetic erosion is challenging (Mallet, 2007), and modeling ancient reticulation requires large datasets and sophisticated analyses. High-throughput sequencing has made these investigations feasible (Dodsworth et al., 2019). Target enrichment methods such as Hyb-Seq enable the analysis of thousands of low-copy nuclear loci (Karbstein et al., 2021), whereas genome skimming can recover plastid genomes (tracing maternal inheritance), ribosomal DNA (biparental nuclear signal), and repetitive elements such as satellites and transposable elements (Dodsworth, 2015; Cavallini et al., 2019). Repeat abundance and sequence similarity offer complementary, alignment-free tools for phylogenetic reconstruction (Dodsworth et al., 2019; Vitales et al., 2020).

The Malvatheca clade (Malvaceae), which originated near the K–Pg boundary (∼66 Mya; Hernández-Gutiérrez & Magallón, 2019; Cvetković et al., 2021), serves as a system for investigating these processes. It comprises Bombacoideae (17 genera, ∼160 species), Malvoideae (78 genera, ∼1,670 species), and the recently recognized Matisioideae (three genera, ∼138 species) (Baum et al., 2004; Carvalho-Sobrinho et al., 2016; Cvetković et al., 2021; Colli-Silva et al., 2025). Matisioideae, formerly the tribe Matisieae, was elevated to subfamily rank by Colli-Silva et al. (2025) after consistent recovery of its monophyly, despite its morphological distinctiveness. While Matisioideae is a well-supported lineage, its position—sister to Bombacoideae or to a recircumscribed Malvoideae—remains poorly resolved, even with phylogenomic data. Malvoideae includes widely cultivated herbs such as cotton, hibiscus, and okra (Baum et al., 2004), whereas Bombacoideae consists mainly of tropical trees, including ecologically and culturally significant species such as the baobab (*Adansonia* L. sp.) and the kapok tree (*Ceiba pentandra* (L.) Gaertn.) (Wickens et al., 2008; Zidar et al., 2009). Several genera, such as *Ochroma* Sw. and *Chiranthodendron* Larreat., placement remain uncertain due to conflicting morphological and molecular evidence. This phylogenetic uncertainty, coupled with the clade’s history of polyploidy, makes Malvatheca a valuable system for studying reticulate evolution in ancient lineages.

The subfamilies within Malvatheca appear to have followed contrasting genomic trajectories shaped by postpolyploid diploidization. Bombacoideae is predominantly composed of species with high chromosome numbers (2n = 86–276; Costa et al., 2017), whereas Malvoideae ranges from 2n = 10–130, with a modal value near 2n = 16 (Tate et al., 2005). Genomic studies have reported signals of reticulate allopolyploidization in Malvatheca and his consequences to their unresolved phylogentic relationships (Hernández-Gutiérrez et al., 2022; Sun et al., 2024; Yang et al., 2025; Zhang et al., 2025). To address this question, we analyze ribosomal DNA, plastome and repeatome data of 22 Bombacoideae, 32 Malvoideae, 2 Matisioideae, and 6 *incertae sedis* taxa under a comparative phylogenomic framework. Specifically, we ask the following questions: (i) Is there evidence of reticulate evolution in the group, and is it associated with taxa of uncertain placement? (ii) Are there repeatome signatures linked to ancient WGDs? (iii) Does lifeform correlate with the distinct diploidization pathways observed in Bombacoideae and Malvoideae?

## Materials and Methods

### Data acquisition and repeatome characterization

We analyzed 62 species of the Malvatheca clade, including 32 Malvoideae, 22 Bombacoideae, 2 Matisioideae, and 6 *incertae sedis*. Two *Theobroma* species (Bytnerioideae) were included as outgroups. Species names and NCBI accession codes (when available) are listed in Table 1. Target-capture sequencing was used to expand the representation of Bombacoideae, following Costa et al. (2021).

**Table 1.** Species and accessions used.

Repeatome analyses were performed with the RepeatExplorer2 pipeline implemented in the Galaxy server (https://repeatexplorer-elixir.cerit-sc.cz; Novák et al., 2020), which clusters reads on the basis of sequence similarity. Two approaches were applied: (i) individual analyses, where each species was analyzed separately to characterize its repetitive fraction, and (ii) comparative analysis, where concatenated reads from all species were analyzed jointly to compare repeat composition across lineages.

For individual analyses, ∼0.1× genome coverage per species was used (Table 1). For species with unknown genome sizes, values were estimated on the basis of closely related taxa. For the comparative analysis, reads were normalized to ∼0.05× per species, yielding a total of 61,288,798 reads. In both cases, clustering was performed using 90% sequence similarity and 55% minimum overlap. The proportions of repeat lineages were calculated as the number of reads per cluster relative to the total number of reads analyzed, excluding chloroplast and mitochondrial sequences identified as potential contaminants.

### Correlations between karyotypic, genomic and ecological traits

Following Schley et al. (2022), we treated the genome as a “community” and repetitive element lineages as “species,” allowing the application of community ecology metrics to genome composition. Repeat diversity was quantified via the Shannon diversity index (H; Shannon, 1948), which measures the probability that two randomly chosen repeat copies belong to the same lineage. Higher H values indicate greater repeat diversity.

For each species, diversity indices were calculated from repeat abundances obtained from the RepeatExplorer individual analyses. To test whether repeat diversity was associated with chromosome number, we performed a Pearson correlation between log₁₀-transformed Shannon index values and log chromosome counts across Malvatheca species via the stats package in R (R Core Team, 2019).

To further examine potential ecological correlates, we compiled chromosome numbers and growth habits (herbaceous vs. woody) for 84 species from the literature. Relationships between habit and chromosome number were visualized with boxplots constructed in PAST 4 (Hammer et al., 2001).

### Plastome, nuclear and repeat-based phylogenies and reticulate evolution

Plastomes and ribosomal DNA (rDNA) were assembled for all the sampled species via the complete plastome of *Theobroma cacao* (NC_014676.2) and the rDNA sequence JQ228369.1 as a reference. The reads were mapped against these references in Geneious v6.0.3 (Kearse et al., 2012) via the “Map to reference” function. The consensus plastomes were aligned with MAFFT (Katoh & Standley, 2013). Maximum likelihood (ML) phylogenies were inferred with 1,000 bootstrap replicates in Geneious v9.1.8 via the FastTree plugin (Price et al., 2009). The resulting trees were visualized and edited in FigTree (http://tree.bio.ed.ac.uk/software/figtree/).

Phylogenetic inference on the basis of repeat abundances was performed following Dodsworth et al. (2015). The abundances of repeat classes obtained from comparative repeatome analyses were treated as quantitative characteristics. Parsimony analyses were conducted in PAST 4 (Hammer et al., 2001).

Phylogenetic networks were reconstructed to evaluate RE. A neighbor-joining tree was first inferred from plastid and rDNA alignments via the Tamura–Nei distance model with 10,000 bootstrap replicates. These tree datasets served as the basis for a SplitsNetwork in SplitsTree v6.4.17 (Huson & Bryant, 2006), generated with the Consensus Splits method and the ConsensusOutlier algorithm, with edge weights proportional to counts. To further detect and visualize putative reticulation events, the autumn algorithm was applied within SplitsTree, and the ML nuclear topology was compared with the plastid tree.

## Results

### Comparison between plastidial and nuclear topologies

The plastome dataset (220,387 bp; 15.1% informative sites) produced a well-supported topology (posterior probability > 0.75 at most nodes), recovering three main clades (Supplementary Fig. 1). Clade I (Bombacoideae Maximum Likelihood bootstrap (BS) = 100) included *Ceiba* Mill., *Spirotheca* Ulbr., and *Pseudobombax* Dugand together with *Adansonia*, who are sisters to *Pachira* Aubl. + *Eriotheca* Schott & Endl. *Fremontodendron mexicanum* Davidson. was resolved as the earliest-diverging lineage. Clade II (Malvoideae) was well supported (BS= 100), with *Pentaplaris* L.O.Williams & Standl. as the earliest diverging species and included the incertae sedis genera *Camptostemon* Mast. and *Howittia* F.Muell. Clade III grouped *Quararibea* Aubl. (Matisioideae) with Malvoideae taxa (*Hampea* Schltdl., *Napaea* L.) and *incertae sedis* (*Lagunaria* (A.DC.) Rchb., *Chiranthodendron*, *Ochroma*). Most genera were consistently monophyletic (*Adansonia*, *Ceiba*, *Pseudobombax*, *Bombax* L., *Malva* Tourn. ex L., *Gossypium* L., *Abutilon* Mill., *Sida* L., *Kokia* Lewton), with the exceptions of *Pavonia* Ruiz. and *Hibiscus* L.

The rDNA dataset (6,456 bp; 16.6% informative sites) also recovered clades I–III but revealed several topological conflicts relative to plastomes (Supplementary Fig. 1). Within Bombacoideae, *Fremontodendron* shifted position, whereas *Phragmotheca mammosa* W.S. Alverson (Matisioideae) grouped closer to clade III rather than with Bombacoideae (see Fig. 1). In Malvoideae, *Pavonia schiedeana* Steud. showed the strongest incongruence, moving from clade II in the plastid tree to clade III in the nuclear tree. Additional conflicts were observed in the placement of *Bombax ceiba* L., *Scleronema micranthum* (Ducke) Ducke, and the groupings *Gossypium* + *Kokia* and *Decaschistia* Wight & Arn. + *Hibiscus* + *Alyogyne* Alef.

**Fig. 1.**
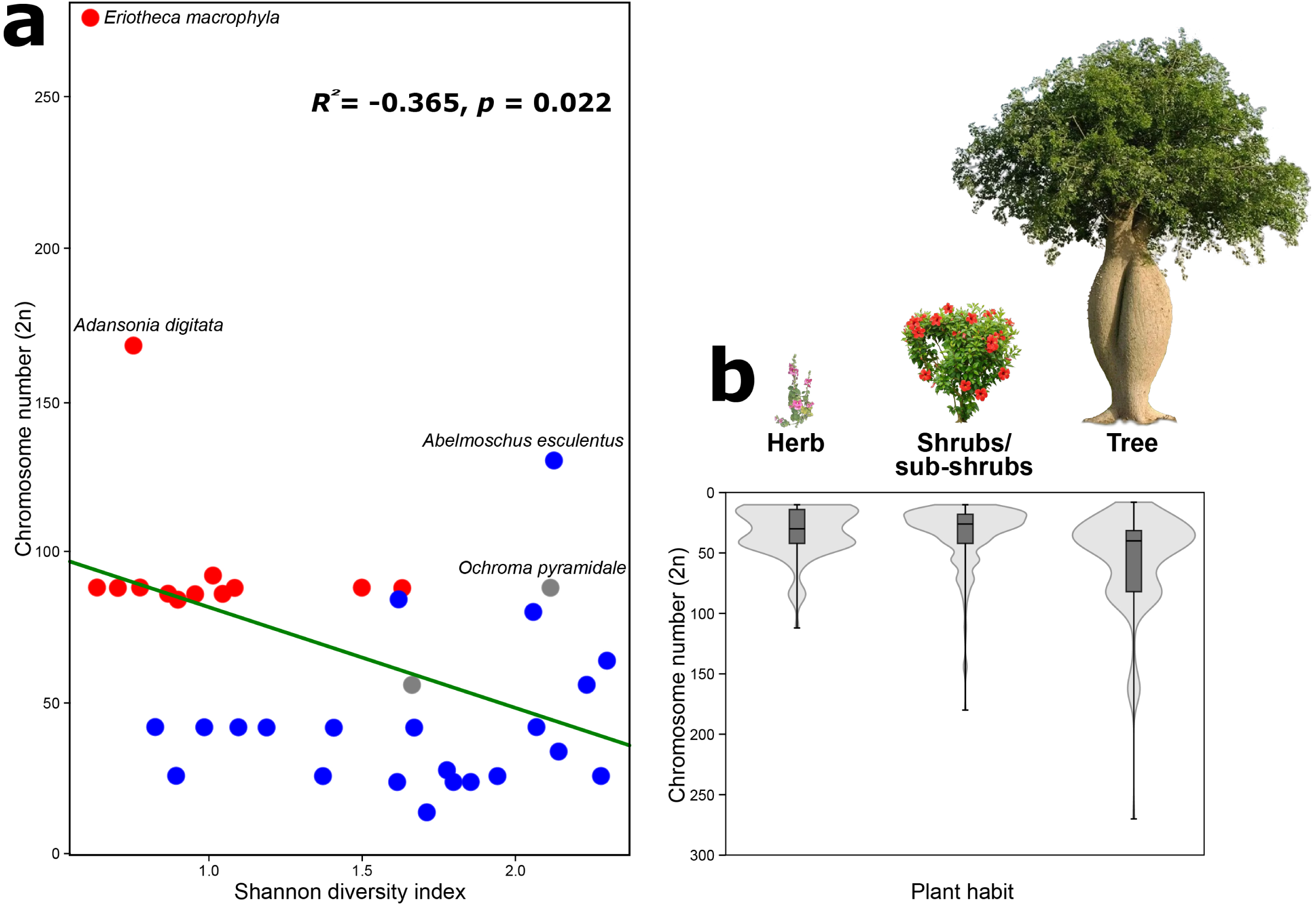
Phylogenetic relationships within the Malvatheca clade inferred from plastome (left) and nuclear rDNA (right) sequence data. Tip labels are color-coded by taxonomic group: Malvoideae (blue), Bombacoideae (red), Matisioideae (green), and incertae sedis (gray). Lines connect the same taxa across plastid and nuclear topologies, highlighting congruences and conflicts.

### Reticulate evolution in Malvatheca

We tested for reticulate evolution (RE) via two complementary approaches in SplitsTree: (i) consensus networks for plastome and rDNA datasets analyzed separately and (ii) an autumn algorithm to reconcile incongruences between plastid and nuclear trees (Fig. 2). Consensus networks revealed extensive phylogenetic conflict, particularly in the rDNA dataset (Fig. 2a). In both plastome- and rDNA-based networks, reticulation was concentrated at the backbone of clade III, especially near taxa currently classified as incertae sedis. Additional reticulations were detected in Bombacoideae (within *Adansonia* and the *Bombax* + *Eriotheca* + *Pachira* + *Spirotheca* clade) and in Malvoideae, where conflict centered on a trifurcation involving *Lawrencia* Hook., the *Malva–Althaea* L.*–Lavatera* L. group, and a *Sida–Abutilon* lineage. The plastome dataset also showed reticulation at the base of *Sida* + *Abutilon*.

**Fig. 2.**
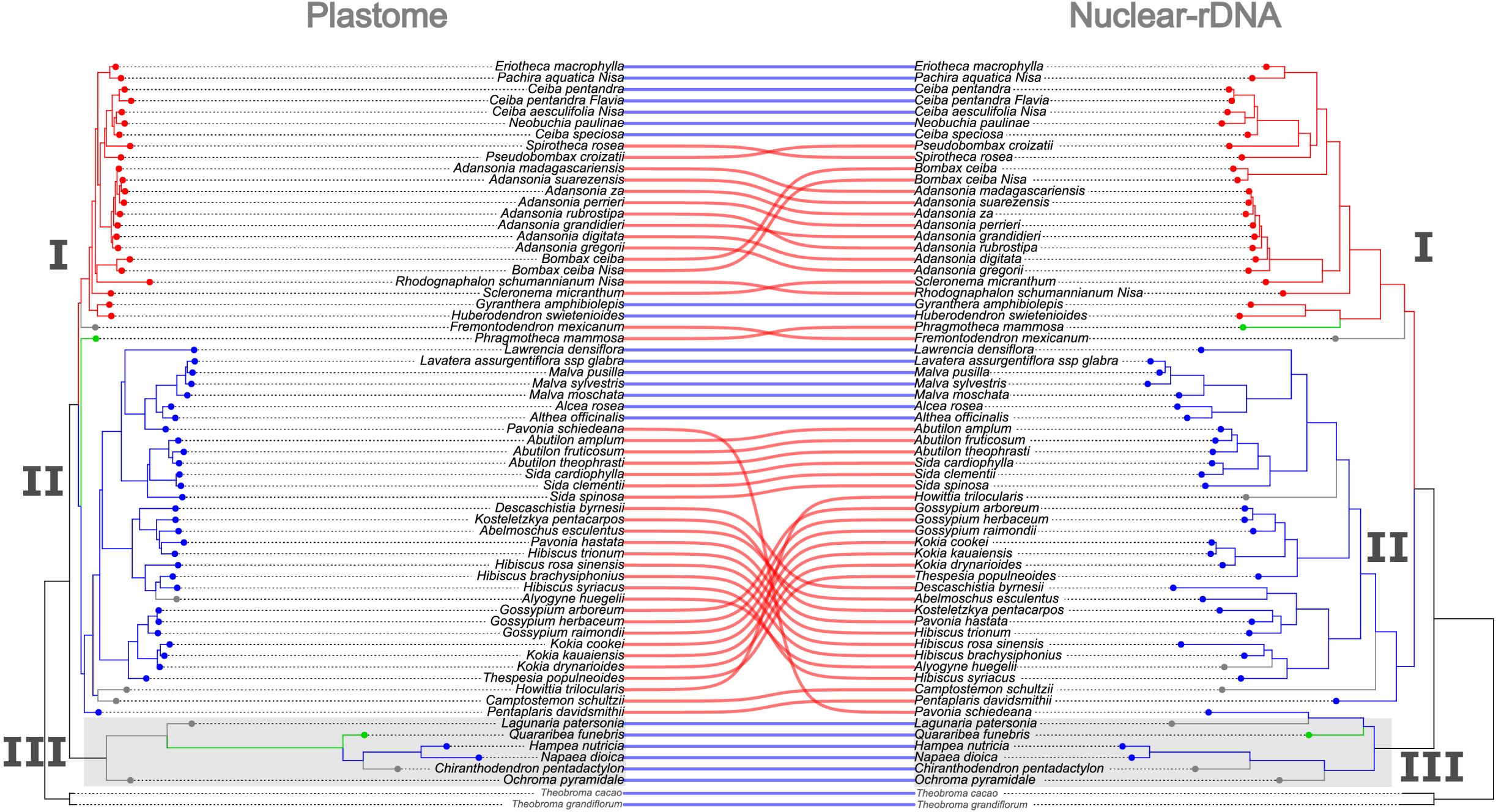
Phylogenetic conflict and reticulation in the Malvatheca clade (Malvaceae). (a) NeighborNet consensus networks for plastome and rDNA datasets, with orange edges highlighting reticulation. (b) Reticulate phylogeny from the autumn algorithm, showing 17 modeled hybridization events (orange connections) between the plastome and rDNA trees. Tip labels are color-coded: Malvoideae (blue), Bombacoideae (red), Matisioideae (green), and incertae sedis (gray). In (a) bold species highlight incertae sedis, while in (b) bold species have been illustrated alongside the phylogenetic tree.

The autumn algorithm identified 17 explicit reticulation events linking the plastome and rDNA topologies (Fig. 2b). Within clade III, one event connected *Quararibea funebris* (La Llave) Vischer and *Ochroma pyramidale* (Cav. ex Lam.) Urb., giving rise to the *Chiranthodendron* + *Hampea* + *Napaea* lineage. Eight events occurred within Malvoideae, including reticulations involving *Pavonia schiedeana* Steud. and *Howittia trilocularis* F.Muell. The remaining eight were detected within Bombacoideae, three of which are inside *Adansonia*. One cross-clade event connected the base of clade II with the *Huberodendron* Ducke + *Gyranthera* Pittier lineage, which was ultimately associated with the Matisioideae genus *Phragmotheca*.

### Repeatome diversity in the Malvatheca clade

In the individual repeatome analyses, the sequencing depth ranged from 40,572 reads in *Bombax ceiba* (accession 2) to 2,633,822 in *Hibiscus syriacus*. The proportion of repetitive DNA varied widely, from 4.8% in *Pseudobombax croizatii* A. Robyns to 56.4% in *Spirotheca rosea* (Seem.) P.E.Gibbs & W.S. Alverson (Bombacoideae), and from 3.4% in *Howittia trilocularis* to 65.4% in *Abutilon amplum* Benth. (Malvoideae) (Fig. 3).

**Fig. 3.**
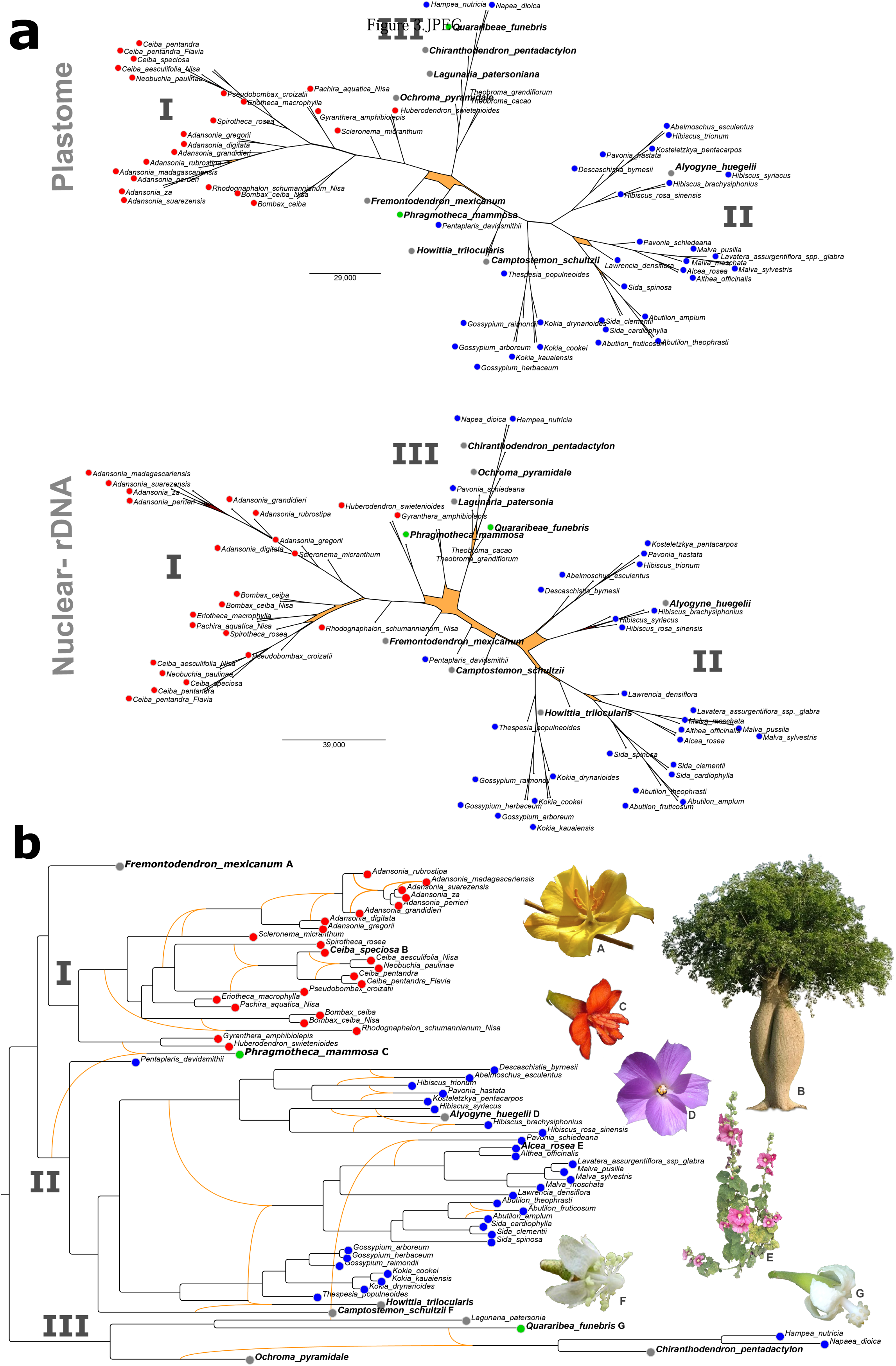
Repeatome composition of the Malvatheca clade. The plastome-based phylogeny (left) is shown alongside the repeat profiles for each species (right). Stacked bars indicate the relative proportions of major repeat classes (normalized to 100%), whereas dots represent their estimated absolute genomic abundance (Mb).

Both subfamilies were dominated by Class I transposable elements, particularly LTR retrotransposons of the *Ty3-type* superfamily. Their abundance ranged from 0.5% in *Malva sylvestris* L. to nearly 50% in *Ceiba speciosa* (A.St.-Hil., A.Juss. & Cambess.) Ravenna. Within Bombacoideae, Tekay elements predominated across four genera, whereas Ogre elements also occurred at moderate to high levels. In Malvoideae, Tekay and Ogre were generally present in moderate proportions, with a few species showing elevated abundances of Athila elements. Among *Ty1-copia* elements, Angela and SIRE were the most widespread across both subfamilies, with SIRE being particularly enriched in *Sida cardiophylla* Domin. and one *Bombax ceiba* accession (Fig. 3; Supplementary Material). The satellite DNA content also varied considerably, from 0.02% in *Gossypium arboreum* L. to 33.9% in *Adansonia grandidieri* Baill. Bombacoideae, especially *Adansonia*, tended to have relatively high satellite DNA abundances (4.9–33.8%). Despite this variation, no genomic synapomorphies based on repeat presence/absence were detected that clearly distinguished Bombacoideae from Malvoideae (Fig. 3).

Comparative repeatome analysis revealed limited sharing of repetitive DNA clusters between subfamilies. Within Bombacoideae, intrageneric similarity was high: *Adansonia* species shared the most repeat clusters, and *Pseudobombax croizatii* and *Pachira aquatica* Aubl. Both genera presented elevated proportions of *Ty3-type* Ogre chickens and CRMs. In contrast, Malvoideae species presented highly divergent repeatomes, with even congeneric taxa (e.g., *Malva sylvestris* and *M. moschata* L.) sharing few clusters (Supplementary Fig. 2).

The repeat-based phylogeny recovered three major clades but with generally weak support (Supplementary Fig. 2). Clade III showed extensive mixing, grouping representatives of all subfamilies without resolution. Several genera were not recovered as monophyletic, including *Adansonia* (Bombacoideae) and *Abutilon*, *Sida*, *Hibiscus*, and *Malva* (Malvoideae). Three Bombacoideae species (*Phragmotheca mammosa*, *Gyranthera amphibiolepis* W. Palacios, *Huberodendron swietenioides* (Gleason) Ducke) and four Malvoideae species (*Camptostemon schultzii* Mast., *Pentaplaris davidsmithii* Dorr & C. Bayer, *Alyogyne huegelii* (Endl.) Fryxell, *Pavonia schiedeana* Steud.) shifted into clade III. Additional rearrangements, such as altered relationships among *Adansonia*, *Scleronema* Benth, and *Rhodognaphalon* (Ulbr.) Roberty (clade I) and the placement of *Pavonia*, *Decaschistia*, and two *Hibiscus* species in clade II (Supplementary Fig. 3).

Comparison with the rDNA tree revealed similar patterns: multiple species from clades I and II were grouped into clade III, with further positional changes within clades (e.g., *Bombax ceiba*, *Fremontodendron*, and *Rhodognaphalon* in clade I; *Howittia*, *Hibiscus*, and *Decaschistia byrnesii* Fryxell in clade II) (Supplementary Fig. 3).

### Diversity of repeats and correlation with chromosome number

The repeat diversity of Malvatheca, as measured by the Shannon (H) and Simpson (D) indices, ranged from very low values in *Rhodognaphalon schumannianum* (A. Robyns) A. Robyns (H = 0.4, D = 0.17) to high values in *Hibiscus brachysiphonius* F. Muell. (H = 2.4, D = 0.9). Malvoideae species generally presented greater repeat diversity (H = 0.8–2.4; D = 0.3–0.9) than did Bombacoideae, which tended toward lower values (Fig. 4a). Chromosome number was negatively correlated with repeat diversity (Shannon: R = –0.36, p = 0.022; Simpson: R = –0.39, p = 0.011), indicating that species with higher chromosome counts, such as Bombacoideae, typically presented fewer diverse repeatomes, whereas Malvoideae (lower 2n) presented greater diversity (Fig. 4b).

**Fig. 4.**
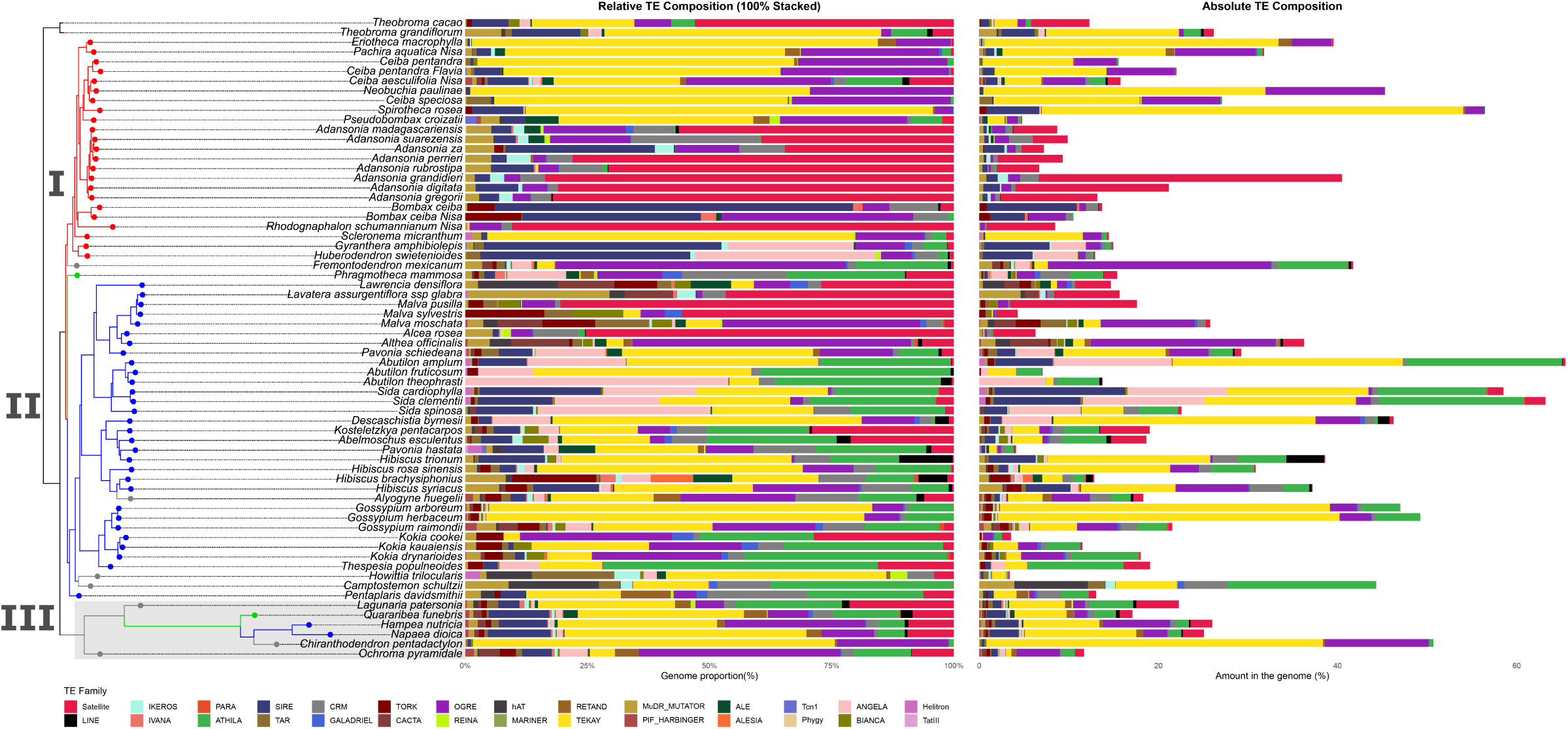
Repeat diversity and chromosome number variation in Malvaceae. (a) Shannon diversity index of repetitive DNA elements in Malvatheca species, colored by taxonomic group: Bombacoideae (red), Malvoideae (blue), and incertae sedis (gray). (b) Chromosome number distributions across growth habits (herbs, shrubs/subshrubs, and trees) compiled from Malvaceae species.

To assess whether chromosome number is correlated with growth habits, we compiled data from 84 Malvaceae species. Trees presented consistently greater chromosome numbers than shrubs and herbs did (range: 8–328; median = 30), supporting an association between the arboreal habit and large chromosome complements (Fig. 4b).

## Discussion

### Would the diversification of Malvatheca be driven by reticulate evolution?

Our data provide phylogenetic evidence for RE in the Malvatheca clade, corroborating previous genomic analyses (Karimi et al., 2020; Wan et al., 2024; Zhang et al., 2025). These reticulation events appear to have occurred on different timescales (Hernández-Gutiérrez et al., 2022; Yang et al., 2025; Stull et al., 2023), ranging from more recent lineages (e.g., the 21.6 Mya genus *Adansonia*; Wan et al., 2024) to the 53.5 Mya origin of Bombacoideae (Zizka et al., 2020; Hernández-Gutiérrez et al., 2022). Hybridization has long been recognized as a fundamental evolutionary force shaping plant diversity, with impacts ranging from immediate reproductive isolation to long-term adaptive radiation (Rieseberg, 1997; Soltis et al., 2004; Chester et al., 2012). One of the main signals in our analyses is the recovery of a well-supported clade III, which groups taxa previously considered incertae sedis together with representatives of Matisioideae and Malvoideae. This clade is consistently associated with reticulation points in both plastid and nuclear datasets, suggesting a complex hybrid origin (Hernández-Gutiérrez et al., 2022; Yang et al., 2025). The morphological and cytogenetic heterogeneity of its members supports this view. Importantly, our sampling does not allow us to test the monophyly of the subfamily Matisioideae, which was recently established in broader phylogenomic analyses with a strong taxonomic delimitation focus (Colli-Silva et al., 2025). Instead, the shifting placement of taxa such as *Phragmotheca* likely reflects reticulation and/or methodological limitations (e.g., low coverage).

While nuclear and plastid datasets provided strong phylogenetic resolution for Malvatheca, the repeat-based approach (Dodsworth et al., 2015) failed to resolve relationships. This loss of signal may reflect (i) the impact of reticulation, since hybridization can alter the presence, absence, or abundance of repeats (Parisod et al., 2009), and/or (ii) the deep divergence of Bombacoideae (53.5 Mya; Zizka et al., 2020). Because repeats undergo rapid evolutionary turnover, their phylogenetic utility can erode in ancient groups. Nevertheless, recent studies have shown that repeatome composition can retain signals and even capture ancient hybridization through characteristic proliferation patterns (Castro et al., 2024; Hlavatá et al., 2024). The contrast across plant groups highlights the lineage- and timescale-specific nature of repeat data. For example, in *Erythrostemon* Klotzsch (Leguminosae, 33.6 Mya), systematic variation in repeat composition revealed discordant profiles consistent with ancient origins (Castro et al., 2024). Similarly, in *Amomum* L. (Zingiberaceae, 19.3 Mya), bursts of repeat proliferation coincided with inferred hybridization events, driving genome size changes that track clade divergence (Li et al., 2023; Hlavatá et al., 2024).

Genomic-scale phylogenies further demonstrated that reticulate evolution is pervasive across Malvaceae, challenging tree-based models (Hernández-Gutiérrez et al., 2022; Yang et al 2025; Zhang et al., 2025). Using 268 nuclear loci across 96 genera, Hernández-Gutiérrez & Magallón. (2019) detected extensive discordance attributable to incomplete lineage sorting and introgression, showing that bifurcating trees fail to capture the family’s evolutionary complexity. Reticulation manifests at multiple scales, from recent interspecific gene flow to ancient allopolyploid events. Hyb-Seq datasets and network inference have revealed well-supported introgression in baobabs (*Adansonia*), clarifying floral homoplasy and pollination biology (Karimi et al., 2020). Similarly, independent hybridization and introgression across *Gossypium* highlight the role of allopolyploidy in diversification (Zhang et al., 2025). Cytonuclear processes in *Hibiscus* (Pfeil, 2002) involve chloroplast capture, whereas cytogenetic work in *Eriotheca* (Serra et al., 2022) confirms that hybridization and polyploidy remain active processes. Together, these findings underscore that phylogenetic networks, rather than bifurcated trees, more accurately represent Malvaceae evolution and highlight the central role of RE in shaping its genomic and phenotypic diversity.

### The repeatome compositions of the Malvatheca clade subfamilies

Genomic studies indicate that the Malvatheca clade has undergone multiple rounds of polyploidization, including reticulate allopolyploidy and subsequent dysploidy, which have played central roles in its diversification (Hernández-Gutiérrez et al., 2022; Yang et al., 2025; Zhang et al., 2025). Recent chromosome-scale assemblies of *Adansonia* spp., *Bombax ceiba* and *Ceiba pentandra* have provided unprecedented insights into genome structure and repeat composition across Malvaceae (Wan et al., 2024; Shao et al., 2024). Complementary phylogenomic studies, such as those on *Hibiscus* L., have revealed at least three independent whole-genome duplication (WGD) events (Eriksson et al., 2021), highlighting the recurrent nature of polyploidy within the family. These duplications supplied raw genetic material for evolutionary novelty, whereas subsequent post-polyploid diploidization processes reshaped karyotypes and contributed to current genomic diversity patterns.

The broader evolutionary significance of WGD has been well documented across angiosperms. Large-scale comparative studies have shown that WGDs are not randomly distributed in time but cluster during episodes of environmental stress (Van de Peer et al., 2017; Teng et al., 2022). This has given rise to the so-called “polyploidy paradox”: although WGD frequently occurs, only a subset of polyploid lineages survive in the long term (Van de Peer et al., 2021). Several of these surviving events appear to coincide with the Cretaceous–Paleogene (K–Pg) boundary (∼66 Mya), a period of mass extinction. WGDs at this time are hypothesized to have facilitated lineage survival and diversification by buffering against genomic stress and providing redundancy for adaptive innovation (Fawcett et al., 2009; Baduel et al., 2018; Bomblies, 2020). The “polyploid hop” model further proposes that polyploidy creates both challenges and opportunities: initial barriers to establishment may be overcome by long-term adaptive potential (Baduel et al., 2018). Recent studies also suggest that specific life-history traits, such as seed biology, may interact with polyploid status to influence which lineages persist across the K–Pg boundary (Cai et al., 2019; Berry & Jaganathan, 2022).

Within Malvaceae, repeatome dynamics provide important clues to post-WGD genome evolution. Chromosome-scale assemblies have shown that transposable element (TE) proliferation and chromosome restructuring are major forces shaping genome size and karyotype variation (Li et al., 2024; Shao et al., 2024). In Malvatheca, distinct repeatome profiles are observed among subfamilies: Bombacoideae tends to exhibit lower repeat diversity, Malvoideae has higher abundance and diversity, and Matisioideae has intermediate patterns. These differences likely reflect contrasting evolutionary strategies for restructuring genomes after polyploidy, with Bombacoideae favoring karyotypic stability through TE amplification, Malvoideae accumulating a more diverse repeat fraction, and Matisioideae retaining a mixed profile.

Taken together, these patterns suggest that the interplay among ancient WGDs, repeatome evolution, and post-polyploid diploidization has been a major driver of lineage-specific diversification in Malvatheca. Variation in repeat composition and chromosome number across subfamilies reflects different genomic strategies for managing the legacy of polyploidy, highlighting the central role of repetitive DNA in shaping both the stability and flexibility of plant genomes over deep evolutionary timescales.

### The habit correlated with chromosome number

Our results revealed that tree-dominated lineages presented the highest chromosome numbers. In Bombacoideae, striking examples include *Pseudobombax* and *Pachira* (2n = 88–92), *Adansonia digitata* (2n = 160), and *Eriotheca* species, with exceptionally elevated counts of 2n = 194–276 (Costa et al., 2017). A similar pattern was described for incertae sedis, such as *Ochroma pyramidale* (2n = 84) (Costa et al. 2017). A similar trend occurs in the exclusively arboreal Tilioideae, where counts range from 2n = 82 to 2n = 328 (Pigott, 2002; 2012). These data reflect the trend toward ancient whole-genome duplication (WGD) events, followed by the lineage-specific conservation of high chromosome numbers in long-life-cycle species.

Shrubby lineages, such as *Hibiscus* (2n ≈ 52) and *Pavonia* (2n ≈ 56), present intermediate values, possibly representing transitional stages between herbaceous and woody forms. In contrast, herbaceous genera such as *Sida* are chromosomally conserved, typically ranging from 2n = 14–32 (Skovsted, 1935; Fernández et al., 2003). These reductions likely result from dysploidy — the step-wise loss or fusion of chromosomes — a process often associated with short-lived, fast-reproducing life cycles (Siljak-Yakovlev et al. 2017).

The habit–chromosome relationship thus reflects contrasting evolutionary pressures. In woody clades such as Bombacoideae, ancient polyploidization established high baselines that were further expanded by more recent polyploidy (Marinho et al., 2014). Herbaceous lineages, in turn, underwent systematic reductions, which is consistent with patterns observed in other families, such as Asteraceae, where perennial-to-annual transitions are linked to smaller karyotypes (Watanabe et al., 1999). Selective pressures on herbaceous species — favoring rapid cell cycles and short generation times — likely reinforced this reduction. Similar correlations between life form and karyotype have been documented across angiosperms (e.g., Malpighiaceae, Lombello & Forni-Martins, 2003; Ehrendorfer, 1989), but Malvaceae provides one of the clearest examples: Bombacoideae, dominated by trees, consistently carries higher chromosome numbers than Malvoideae, which include mostly herbs and shrubs. These results suggest that major life form transitions are accompanied by fundamental changes in genome organization.

## Conclusions

Our phylogenomic analyses provide strong evidence for reticulate evolution in the Malvatheca clade (Malvaceae), underscoring its potential role as a driver of diversification in the family. Ancient hybridization events, coupled with allopolyploidy and subsequent diploidization, have left lasting imprints on the genomic architecture and subfamily relationships. Nuclear and plastid datasets revealed extensive reticulation, whereas repeatome analyses proved less effective for deep phylogenetic resolution, reflecting rapid lineage-specific turnover. This contrast highlights the importance of multilayered genomic approaches for reconstructing complex evolutionary histories.

Comparative genomic and cytogenetic evidence also reveals subfamily specific patterns in karyotype evolution. Tree-dominated lineages, particularly Bombacoideae, exhibit exceptionally high and variable chromosome numbers arising from successive polyploidization, whereas herbaceous taxa display reduced counts through dysploidy and genome downsizing. These correlations suggest a close interplay between life form, polyploidy, and genome structure: woody lineages preserve high baselines established by ancient WGDs, whereas herbaceous taxa evolve toward reduced karyotypes under different life-history constraints.

Taken together, our results demonstrate how hybridization, polyploidy, and repeat dynamics have jointly shaped Malvaceae diversification. This family offers a compelling model for understanding the integration of reticulate processes, structural genome evolution, and life-history strategies in driving morphological and ecological innovation. Future research incorporating chromosome-scale assemblies and functional studies will be critical for disentangling the genomic consequences of WGD and reticulation across this diverse plant lineage.

## Capitons

Supplemetary Fig. 1 Phylogenetic relationships inferred from whole plastome, nuclear and repeat abundance data. Maximum Likelihood trees for the plastome and nuclear genome are shown followed by a Neighbor-Joining tree based on repeat abundance. Support values are shown on nodes for all trees.

Supplementary Fig. 2. Phylogeny and repeatome analysis of the Malvatheca clade (Malvaceae). The plastome-based phylogeny (left) shows the evolutionary relationships between species. The heatmap (right) compares the abundance of different repetitive element classes across these species, with brown indicating high abundance and light colors indicating low abundance.

Supplementary Fig. 3 Phylogenetic relationships within the Malvatheca clade inferred from inferred from plastome and nuclear rDNA (left) and repeats abundance (right) sequence data. Tip labels are color-coded by taxonomic group: Malvoideae (blue), Bombacoideae (red), Matisioideae (green), and *incertae sedis* (gray).

Supplementary Table. 1 Malvatheca repeats amount into genome.

Supplementary Table. 2 Repeat category proportions by species.

Supplementary Table. 3 Number of reads analyzed by species.

Supplementary Table. 4 Repeat abundance matrix used to infer the repeat phylogeny.

Supplementary Table. 5 Correlation between habit and chromosome number.

Supplementary Table. 6 Values of repeat diversity indexes by species.

